# Association between the magnitude of melatonin circadian rhythm and personality in the threespine stickleback

**DOI:** 10.1101/2023.01.30.526243

**Authors:** Morgane Anita Philippe, Nadia Aubin-Horth

**Author notes:** Corresponding author: Nadia Aubin-Horth.

## Abstract

Melatonin secretion follows a circadian pattern with a maximum level at night in many species. However, in zebrafish (*Danio rerio*, a diurnal fish species) large inter-individual variations in daily rhythmicity of melatonin levels are present and are associated with variation in behaviour. Melatonin secretion rhythm of proactive individuals that are more active and exploratory are of larger amplitude compared to reactive individuals. In threespine sticklebacks (*Gasterosteus aculeatus*), a nocturnal species, inter-individual variability of behaviour is well described. However, inter-individual variation of melatonin rhythm and its association with variation in behaviour has never been measured in this species, which would allow to test if patterns found in zebrafish can be generalized for diurnal and nocturnal species. We measured large inter-individual variation in melatonin levels and found that activity was positively correlated with plasma melatonin concentration measured at night. We did not observe any significant difference in nigh-day variation in melatonin concentration between very active and less active groups. However, we found that individuals classified as reactive based on their propensity to wall-hugging, a measure of anxiety in fish, showed large variation in melatonin between night and day, while this rhythm was not seen in proactive individuals that frequently used the centre of the aquarium. Overall, our study suggests that melatonin may directly modulate specific behaviours in wild sticklebacks, and that while interindividual variation in melatonin rhythm may be widespread in fish, different patterns of association with behaviours should be expected.

## INTRODUCTION

Many organisms show daily rhythms of behaviour and physiology (Kumar, 2017). In animals, activity rhythms can be diurnal, nocturnal, crepuscular or arrhythmic depending on the species (Kumar, 2017). These biological rhythms are driven in part by external rhythms in the environment, such as the alternation of the light and dark phases, and photoperiod (duration of the light phase) (Kulczykowska et al., 2010; Kumar, 2017). Other parameters of the environment also play a synchronizing role, such as food intake, temperature and social interactions (Kulczykowska et al., 2010; Kumar, 2017; Zhdanova and Reebs, 2005). Daily rhythms (that have a 24-hour rhythm) are seen for many traits, including sleep cycle, immune system functions, glucose and lipid metabolism, as well as neurotransmitters and hormones secretion (Kumar, 2017). It is generally observed that secretion of melatonin, an hormone, follows a daily pattern with a maximum level at night (Falcón et al., 2011; Kumar, 2017; Pevet et al., 2017). Melatonin targets various tissues and affects their functions (production of neurotransmitters, hormones secretion) (Falcón et al., 2011; Slominski et al., 2012). Melatonin secretion can be characterized as a signal of the dark phase of a 24-hour day, as its downstream effects varies among diurnal and nocturnal species, probably because of different coupling system between light-dark cycle, circadian oscillators and internal functions (Kumar, 2017; Smale et al., 2003). In diurnal species, such as human, melatonin secreted at night inhibits activity and induces somnolence effects (Kumar, 2017). In nocturnal species, such as many rodents, melatonin is also at maximum levels during the dark phase but is instead associated with wakefulness, activity and feeding behaviours (Kennaway, 2019).

While this general model of melatonin secretion rhythm has been validated in different species, it is known that some laboratory mice strains do not secrete melatonin, and that in zebrafish (*Danio rerio*, a diurnal fish species) large inter-individual variations in daily rhythmicity of melatonin levels are present (Kennaway, 2019; Tudorache et al., 2018). In zebrafish, some individuals present the expected large variations between light-phase and dark-phase melatonin levels, while others show no significant change in melatonin levels over 24 hours. When individuals are classified as proactive or reactive based on their risk-taking propensity (measured as the speed to emerge into a new environment), melatonin secretion rhythm of proactive individuals are of larger amplitude compared to reactive individuals (Tudorache et al., 2018). Proactive individuals also show a more pronounced rhythmicity in both activity level and melatonin secretion, i.e. a higher activity level during daytime (measured as average swimming velocities over 30 min) and higher melatonin secretion level at night time compared to reactive individuals. This study did not uncover whether melatonin rhythm and variations in behaviour are linked by a cause-and-effect relationship, or if these two variables are themselves controlled by a third unidentified variable. However, a causal link has been shown in other fish species between melatonin levels and behaviour. Melatonin administration at night decreases locomotor activity and food intake in goldfish (Azpeleta et al., 2010; López-Olmeda et al., 2006) and decreases locomotor activity in European sea bass (Herrero et al., 2007), two diurnal species, while it does not affect locomotor activity in tench (López-Olmeda et al., 2006), a nocturnal species. Furthermore, melatonin exposure attenuates the stress response in the brain in Senegal sole (Gesto et al., 2016; López-Patiño et al., 2013) and in rainbow trout (Conde-Sieira et al., 2014). Finally, in threespine stickleback, melatonin administration affects sexual maturation (Borg and Ekström, 1981; Bornestaf et al., 2001).

Although several species are well known for their inter-individual variations in behaviour (Kight et al., 2013), much less is known about their melatonin secretion rhythm and if there is also inter-individual variation in that phenotype, as seen in zebrafish. Inter-individual variation of melatonin secretion is not well documented, potentially because the majority of studies use laboratory lines that tend to be uniform (Chattoraj et al., 2009; Pevet et al., 2017). It is often assumed that melatonin is secreted with a marked daily rhythm in the same way in most species. But if these rhythms vary between individuals of the same species, we need to better understand this variation, since animal models and their biologically known characteristics are used in different fields, such as in ecotoxicology or medical research. For example, melatonin secretion has been used as a biomarker in ecotoxicology to study the effects of toxic chemicals and other biochemical compounds (Bidwell, 2020). Moreover, melatonin is important for the maintenance of human health (Arendt, 2006; Fishbein et al., 2021; Roenneberg and Merrow, 2016) and the dysregulation of melatonin secretion is involved in different human diseases (Karasek, 2004; Pacchierotti et al., 2001; Pandi-Perumal et al., 2017; Reiter et al., 2007). Melatonin has been also shown to modulate the progression of infections in animal models and in human (Daryani et al., 2018; He et al., 2021; Hu et al., 2017; Silvestri and Rossi, 2013). Finally, having a broader picture of the presence of inter-individual variation in melatonin secretion is a first step before studying whether this variation is the result of genetic variation and if it is under selection in nature. Studying wild species with different characteristics, for example comparing diurnal and nocturnal species, would help to determine the extent of this phenomenon. Furthermore, studying melatonin secretion rhythm in a species that is known for its inter-individual variation in behaviour would also allow to test the association between melatonin rhythm characteristics and behaviour in an additional species to *Danio rerio*.

Our aim was to test the link between melatonin pattern and behaviour variations in wild threespine sticklebacks (*Gasterosteus aculeatus*), whose behaviour can be measured in an automated way. This fish species has been used as a model in behavioural biology (Huntingford and Ruiz-Gomez, 2009; McKinnon et al., 2019; Norton and Gutiérrez, 2019), and in ecotoxicology (Katsiadaki et al., 2007; McKinnon et al., 2019; Ostlund-Nilsson et al., 2006). Personality and inter-individual variability of behaviour, such as exploration, aggressiveness and predator response is well described in threespine sticklebacks (Aubin-Horth et al., 2012; Berger and Aubin-Horth, 2020; Brochu and Aubin-Horth, 2021; Grécias et al., 2018, 2017; Jolles et al., 2019; Lacasse and Aubin-Horth, 2012; Quinn et al., 2012; Talarico et al., 2017). Data from the field and laboratory suggest that threespine sticklebacks are nocturnal. In freshwater, they display a diel vertical migration, being closer to the surface at night than during the day (Quinn et al., 2012). When monitored continuously in the laboratory, they are also more active during the dark phase than during the light phase, although with a high inter-individual variability in activity level (Brochu and Aubin-Horth, 2021). Melatonin has a circadian secretion pattern in threespine stickleback, with a maximum secretion level at night (Kulczykowska et al., 2017). However, inter-individual variation of melatonin rhythm and its association with behaviour has never been measured.

Our hypothesis was that threespine stickleback behaviour and melatonin secretion would vary between individuals. We also hypothesized that a strong rhythmicity of behaviour would be associated with a strong rhythmicity in melatonin secretion. Our predictions were that melatonin levels in plasma and brain measured using an EIA would be high at night and low during the day, as previously shown using HPLC (Kulczykowska et al., 2017) and that melatonin levels would vary between these wild-caught individuals, as seen in that species for behaviours. We also predicted that melatonin levels would be associated with measures of behaviour, as seen in other species. Specifically, based on the observation that sticklebacks are mostly nocturnal but show large inter-individual variation in the phase of activity, and that the most diurnal fish are the most active (Brochu and Aubin-Horth, 2021), we predicted that threespine sticklebacks that exhibit high activity level during the day would have high melatonin levels at night, as seen in a diurnal species with unmanipulated melatonin levels, *Danio rerio* (Tudorache et al., 2018). We also predicted that proactive individuals would present a more contrasted melatonin rhythm (higher melatonin level at night and lower melatonin level during day) than reactive individuals, as seen in *Danio rerio* (Tudorache et al., 2018).

To test these predictions, we first measured exploration and predator response using standard behavioural tests in threespine sticklebacks (Berger and Aubin-Horth, 2020; Grécias et al., 2018, 2017). Note that behaviour tests are usually done during the day in this species (Aubin-Horth et al., 2012; Berger and Aubin-Horth, 2020; Grécias et al., 2018, 2017; Lacasse and Aubin-Horth, 2012; Quinn et al., 2012; Talarico et al., 2017). In summary, exploration is quantified using distance swam and time spent in center of the aquarium; while predator response is quantified by time to freeze, and time spent frozen in response to a predator attack. We defined a proactive and a reactive group of individuals, proactive individuals being more active, risk-taker and exploring novel stimuli more quickly than reactive individuals (Laland et al., 2011; Tudorache et al., 2018). We then measured melatonin in brain and plasma by enzyme immunoassay (EIA) for each individual. Melatonin dosage has already been carried out using whole body measurements in zebrafish (Tudorache et al., 2018) and in plasma of European sea bass (Bayarri et al., 2002), Senegal sole (Bayarri et al., 2004) and Atlantic salmon (Porter et al., 2001; Randall et al., 1995). Melatonin has been measured in the brain of threespine sticklebacks by HPLC (Kulczykowska et al., 2017) and RIA (Sokołowska et al., 2004). One objective of this study was to refine a standard EIA protocol to measure the concentration of melatonin in small samples: the brain and plasma of threespine sticklebacks during day and night. Finally, using these two datasets, we quantified the association of melatonin levels with individual behaviour and with the proactive/ reactive classification.

## MATERIALS AND METHODS

### Fish sampling and housing

We collected threespine sticklebacks (*Gasterosteus aculeatus*) from Lac Témiscouata (Québec, Canada) in June 2018, using a beach seine. Individuals were brought back to the animal facility (Laboratoire de Recherches en Sciences Environnementales et Médicales (LARSEM), Université Laval, Québec, Canada) in insulated coolers. Prior to the experiment, which took place in the summer of 2019, they were held in two 1000L water tanks and fed brine shrimp and nutritious flakes twice a day. Water temperature was set to 14°c and light dark cycle to 12h light: 12h dark with lights on at 9am and lights off at 9pm. These wild individuals thus spent one year in a stable laboratory environment before they were studied.

### Experimental design and sampling

All experimental procedures were approved by the Comité de Protection des Animaux de l’Université Laval (CPAUL certificate # 2018066-1). Fish were transferred to the experimentation room at least 3 days before the start of the experiment and placed in 20 L water tanks in groups of 4 individuals maximum. The photoperiod was 12L:12D with lights on at 9am and off at 9pm. The day before the experiment, individuals were isolated in 2L water tanks without food. On the first day of the experiment (in the morning, see Figure 1), the behaviour of each fish was measured in a 45L test aquarium with white plexiglass-covered walls using standard behaviour test to measure exploration and predator response in threespine stickleback (Berger and Aubin-Horth, 2020; Grécias et al., 2018, 2017). All fishes were re-fed at the end of the behaviour test and then fed twice a day as before.

**Figure 1.**
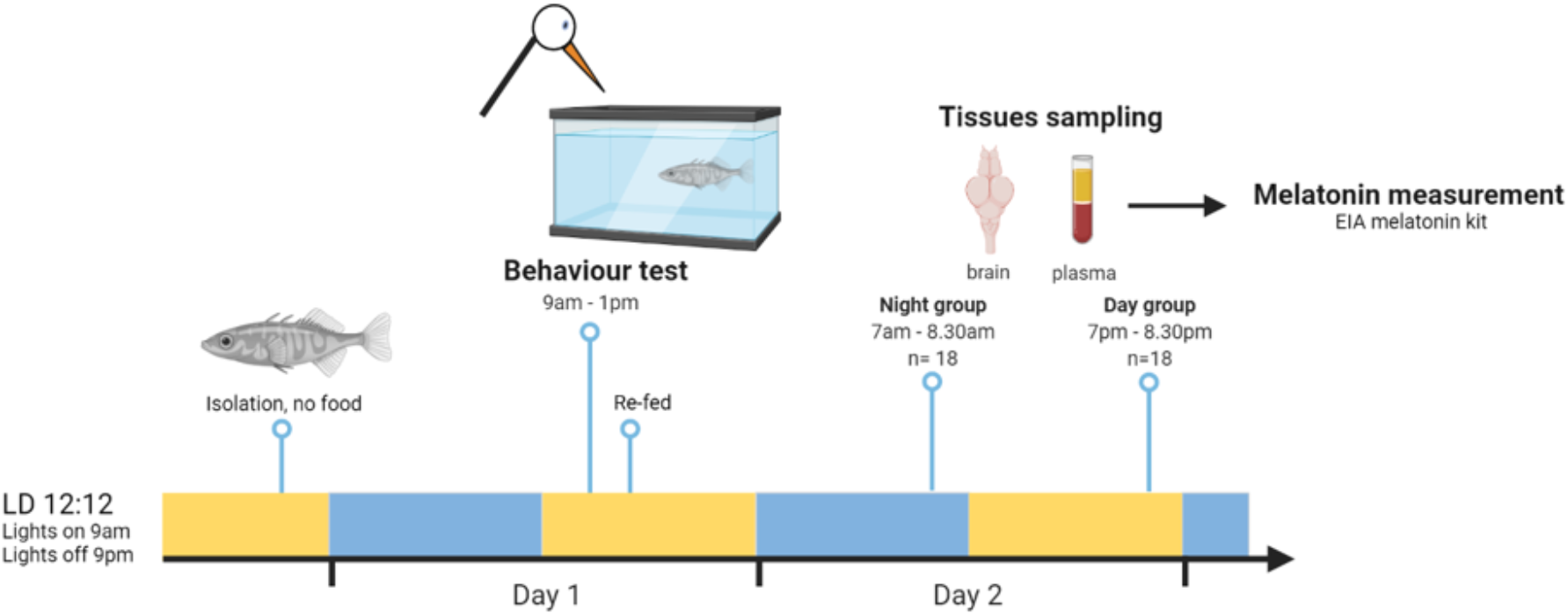
Experimental design used to explore how melatonin rhythm in plasma and brain is linked to personality type in threespine sticklebacks. Sticklebacks behaviours were quantified using standard behaviour test during light phase on day 1. On day 2, plasma and brain tissue were collected during the light or dark phase. Melatonin concentration in plasma and brain samples were determined by EIA.

To measure behaviour, an individual fish was placed in a test aquarium and their behaviour was immediately video recorded during 10 minutes. In this first part of the behaviour test, we measured exploration of a novel environment using travelled distance (“Distance”) and percentage of time spent in the centre of the aquarium (“Time spent in centre”). Then, we simulated a predator attack with a plastic stick and recorded behaviour for another 5 minutes to quantify predator response using the “Time to freeze” and “Time spent frozen” variables. More precisely, we quantified the time taken to freeze after the attack and the time spent frozen until activity was resumed, up to 5 minutes. We tested 39 individuals, 3 of them died before tissues sampling and 2 others showed abnormal behaviour, they were then excluded from the analysis. We thus analysed 34 behaviour test results.

Tissue sampling was performed on the following day. Individuals were randomly divided between a night group and a day group. Tissue sampling was performed during the dark phase for the night group (n=18), between 7 am and 8.30 am, which is the last hour before lights on. We sampled during the light phase for the day group (n=18), between 7 pm and 8.30 pm, just prior to lights off (Figure 1). As we were able to collect tissues on a maximum of 5 individuals between 7.00 and 8.30, we repeated four identical experimental sessions. For each session, a maximum of 10 individuals were tested for behaviour on day 1 and divided in night and day group (5 individuals in each group). Fish were anesthetized by immersion in MS-222 (250 mg L^-1^). Blood samples were collected by tail removal through the caudal vein using heparinized capillary and kept on ice. Fish kept under anaesthesia were finally euthanized by decapitation. Brain tissues were immediately placed in liquid nitrogen and stored at −80°c. Blood samples were centrifuged with a microcentrifuge at 6000rpm for 1 minute or until complete separation from the plasma. Plasma samples were then stored at −20°c.

### Behaviour quantification

Video tracking for behaviour test was performed using an automatic tracking module in EthoVision software (Ethovision XT, Noldus) (Noldus et al., 2001). In the first part of the behaviour test, we measured exploration of a novel environment using travelled distance (“Distance”), time spent being active (“Time active”) and percentage of time spent in the centre of the aquarium (“Time spent in centre”). During the second part of the behaviour test, we quantified the predator response using the “Time to freeze” and “Time spent frozen” in response to a simulated predator attack. We then separated individuals into proactive and reactive groups for each behaviour, according to the median of each behaviour variable. Individuals for whom Distance, Time spent in centre or Time to freeze was greater than the median were considered proactive for that given behaviour, while individuals that had a value of time spent frozen below the median were characterized as proactive. Individuals with the opposite pattern were considered reactive for that behaviour.

### Melatonin measurement in plasma and brain

Melatonin concentration in plasma and brain was determined with a commercial enzyme immunoassay (EIA) kit (IBL, Hamburg, Germany)(Bayarri et al., 2004, 2002). After defrosting, plasma sample volume was completed up to 500μL with washing buffer from the EIA kit. Brain tissue was sonicated in 500μL phosphate-buffered saline (PBS) for 15min using an ultrasonic bath, without heat, then centrifuged for 15min – 15 000g – 4°c and kept on ice. Supernatant was saved for extraction. All samples, standards and controls were purified with C18 extraction columns following manifold vacuum procedure. Briefly, C18 columns were conditioned with 1ml absolute methanol and 1ml bi-distilled water. 500μl of each sample, standard and control were applied on conditioned columns, followed by 700μl bi-distilled water. Columns were washed with 2x 1ml 10% methanol. Elution of extracts was performed with 1ml absolute methanol. Finally, the evaporation of methanol was performed by air drying. Dried extracts were stored at −20°c for a maximum of 24h until EIA test procedure. When ready for EIA test procedure, extracts were reconstituted with 105μl bi-distilled water (instead of 150μl recommended by IBL because of expected low melatonin levels in samples) and vortexed for at least 1min. 50μL of each reconstituted extract were added immediately in duplicate to different wells of an EIA plate precoated with capture antibody. Melatonin concentration was determined according to the standard EIA protocol.

We used a four-parameters logistic model to calculate melatonin concentrations in plasma and brain samples. Plasma and brain samples of two individuals in the night group were used to optimize melatonin measurement with EIA kit. Five plasma samples (four in the day group, one in the night group) with an OD outside the standard curve were excluded. One plasma sample in the day group with a very small volume (1.24 μl) and an aberrant melatonin concentration (1176 pg/ml) was excluded. One brain sample in the night group was lost during manipulation. We performed the final analyses using 28 plasma samples (day n=13, night n=15) and 33 brain samples (day n=18, night n=15).

### Data analysis

Statistical analyses were performed using R software v 4.1.0 (R Core Team, 2018). A Shapiro-Wilk normality test showed that behaviour variables and plasma melatonin concentration were not normally distributed. A Spearman’s correlation test was used to test correlation between behaviour variables and plasma melatonin concentration. A unilateral Wilcoxon test was used to test for differences in plasma melatonin concentration between the day and night groups. A two-way ANOVA was performed to evaluate simultaneously the effect of two grouping variables: Day/Night and Proactive/Reactive, on plasma melatonin concentration. Shapiro-Wilk normality test showed that residuals were not normally distributed. A two-way ANOVA was thus performed with base-10 log transformed plasma melatonin data and untransformed behaviour data. Tukey multiple comparison test was used for the post hoc determination of differences between groups and p-values were adjusted for multiple comparisons.

## DATA AVAILABILITY

The complete dataset is available as supplementary table 1. The R script used for the statistical analysis is available as supplementary material 2.

## RESULTS

### Evaluation of behaviour in threespine sticklebacks

We evaluated five individual behaviours to describe two general responses: exploration (Distance, Time active, and Time spent in centre) and predator response (Time to freeze and Time spent frozen). The five behaviours varied between individuals (Table 1). We then separated individuals into proactive (n=17) and reactive (n=17) groups, according to the median of each behaviour variable. Individuals for whom Distance, Time active, Time spent in centre or Time to freeze was greater than the median were considered proactive for that given behaviour, while individuals that had a value of time spent frozen below the median were characterized as proactive. Individuals with the opposite pattern were considered reactive for that behaviour.

**Table 1.** Behaviours associated with exploration and predator response vary among wild threespine sticklebacks. We measured exploration of a novel environment using travelled distance (“Distance”), percentage of time spent being active (“Time active”), and percentage of time spent in the centre of the aquarium (“Time spent in centre”) for 10 minutes. We then quantified the predator response using the “Time to freeze” and “Time spent frozen” in response to a simulated predator attack. n=34.

|  | Exploration |  |  | Predator response |  |
| --- | --- | --- | --- | --- | --- |
|  | Distance (cm) | Time active (%) | Time spent in centre (%) | Time to freeze (s) | Time spent frozen (s) |
| **Median (min-max)** | 1696<br>(554-4041) | 55<br>(21-87) | 6<br>(0-31) | 3.1<br>(0.2-8.0) | 31.3<br>(4.0-137.6) |

We verified if behaviours were correlated with each other and found that our measures of activity (distance and time spent active) were highly correlated (Spearman correlation, r=0.90 p< 2.2e-16). We did all further analyses with travelled distance only. Time spent in centre, a measure of low anxiety, was not correlated with the other two measures of exploration (Spearman correlation, Distance r = −0.11 p = 0.51; Time active r = 0.01 p = 0.95), and the predator response behaviours were also not correlated (Spearman correlation, r = −0.29, p = 0.09).

### Melatonin concentration in plasma and brain

We measured melatonin concentration in plasma and brain of each individual by EIA. The initial amount of melatonin in samples was in the range of 0-2pg, which required us to adapt the EIA recommended protocol. The concentration of melatonin in the samples was nevertheless close to the lower limit of detection of the EIA kit and some plasma samples that were insufficiently concentrated were excluded (see Materials and Methods). The concentration of melatonin in plasma was variable between the 28 individuals available. In the day group, it ranged between 12.6 pg/ml and 110.2 pg/ml, with a median of 37.4 pg/ml, and varied between 40.0 pg/ml and 188.6 pg/ml in the night group, with a median of 62.0 pg/ml. The mean of the plasma melatonin concentrations in the night group was significantly higher than the mean in the day group (Figure 2, Unilateral Wilcoxon test, p = 0.021. Removing the extreme data point of the night group at 188.6 pg/ml: Unilateral Wilcoxon test, p = 0.034). Melatonin concentration in the brain was less variable between individuals and we did not observe a difference between the day (7.2 – 45.0 pg/g) and night groups (11.1 - 46.6 pg/g) (Unilateral Wilcoxon test, p = 0.16). Previous studies obtained similar melatonin concentrations in brain (Kulczykowska et al., 2017; Sokołowska et al., 2004). Contrary to our study, Kulczykowska et al. observed a significant difference between the concentration of melatonin during the day and at night in the brains of threespine sticklebacks. In this study (Kulczykowska et al., 2017) melatonin was extracted in liquid phase and measured by HPLC, suggesting that our C18 column extraction and/or EIA assay protocol is not suitable for brain samples. Considering the technical limitations of our study, we chose to focus our analysis on the results in plasma.

**Figure 2.**
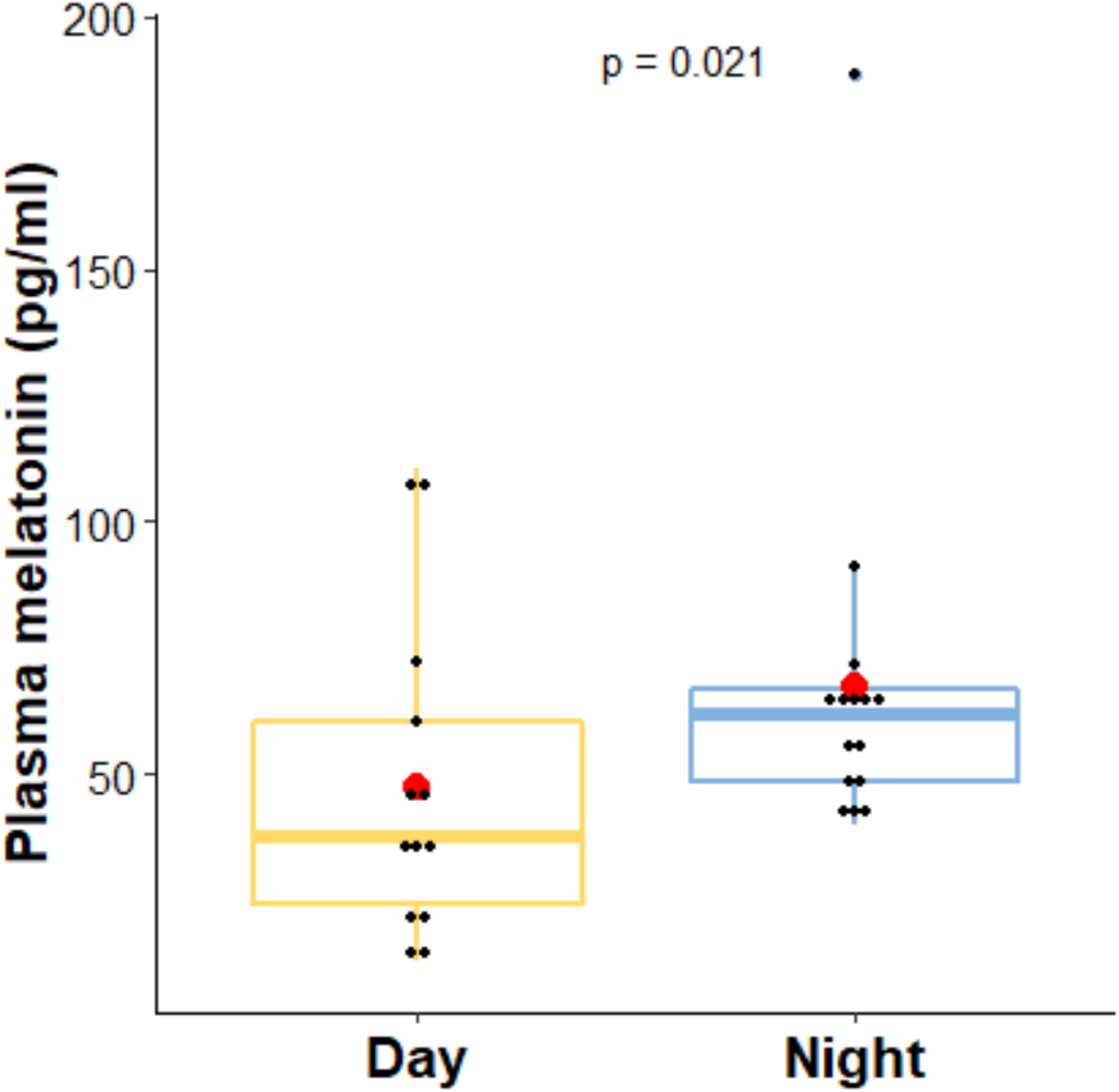
Melatonin concentration in plasma is significantly higher in samples taken at night than during the day. Melatonin concentrations determined by EIA in plasma (pg/ml) of threespine sticklebacks collected during the light phase (n=13) or the dark phase (n=15). The mean of the plasma melatonin concentrations (red points) in the night group (67.5 pg/ml) was significantly higher than the mean in the day group (47.6 pg/ml). Unilateral Wilcoxon test, p = 0.021. Removing the extreme data point of the night group at 188.6 pg/ml: Unilateral Wilcoxon test, p=0.034.

### Correlation between melatonin concentration in plasma and behaviour

Taking all individuals together, we observed a positive and significant correlation between the concentration of plasma melatonin at night and travelled distance (Spearman correlation: r = 0.74, p = 0.002, Table 2). In contrast, melatonin concentration in plasma during the day was not correlated with travelled distance (Table 2). Others behavioural variables (Time spent in centre, Time to freeze, Time spent frozen) were not correlated with melatonin concentration in plasma neither during the day nor at night (Table 2).

**Table 2.** Travelled distance during the day of all individuals is positively correlated with plasma melatonin levels at night. Correlation of behaviour variables and melatonin concentrations determined by EIA in plasma (pg/ml) of threespine sticklebacks collected during the day (n=13) or night (n=15). Other variables are not significantly correlated. Significant correlations are in bold (Spearman’s correlation test).

| Behaviour | Plasma melatonin (pg/ml) |  |
| --- | --- | --- |
|  | Day | Night |
| Distance (cm) | $r = 0.05$<br>$p=0.877$ | $r = 0.74$<br>**$p=0.002$** |
| Time spent in centre (%) | r = 0.44<br>p=0.129 | r = -0.22<br>p=0.411 |
| Time to freeze (s) | r =0.02<br>p=0.949 | r = 0.13<br>p=0.620 |
| Time spent frozen (s) | r = -0.02<br>p=0.934 | r = -0.30<br>p=0.264 |

### Melatonin rhythms in proactive and reactive individuals

We tested if plasma melatonin concentration varied according to time of day (measurement at night or during the day) and to behaviour type, using individuals separated in proactive (n= 17) and reactive (n=17) groups for a given behaviour.

For travelled distance, time of day had a significant effect on melatonin levels (two-way ANOVA, p = 0.032, Table 3). Night time levels of plasma melatonin were higher than during the day (Figure 3). Melatonin levels did not differ significantly according to travelled distance when individuals were divided into high activity and low activity groups, although high activity individuals tended to have higher melatonin levels than low activity ones (Figure 3), (two-way ANOVA, p = 0.056, Table 3). The interaction between Time of Day and Distance was not significant (two-way ANOVA, Distance * Time of day p = 0.847, Table 3).

**Figure 3.**
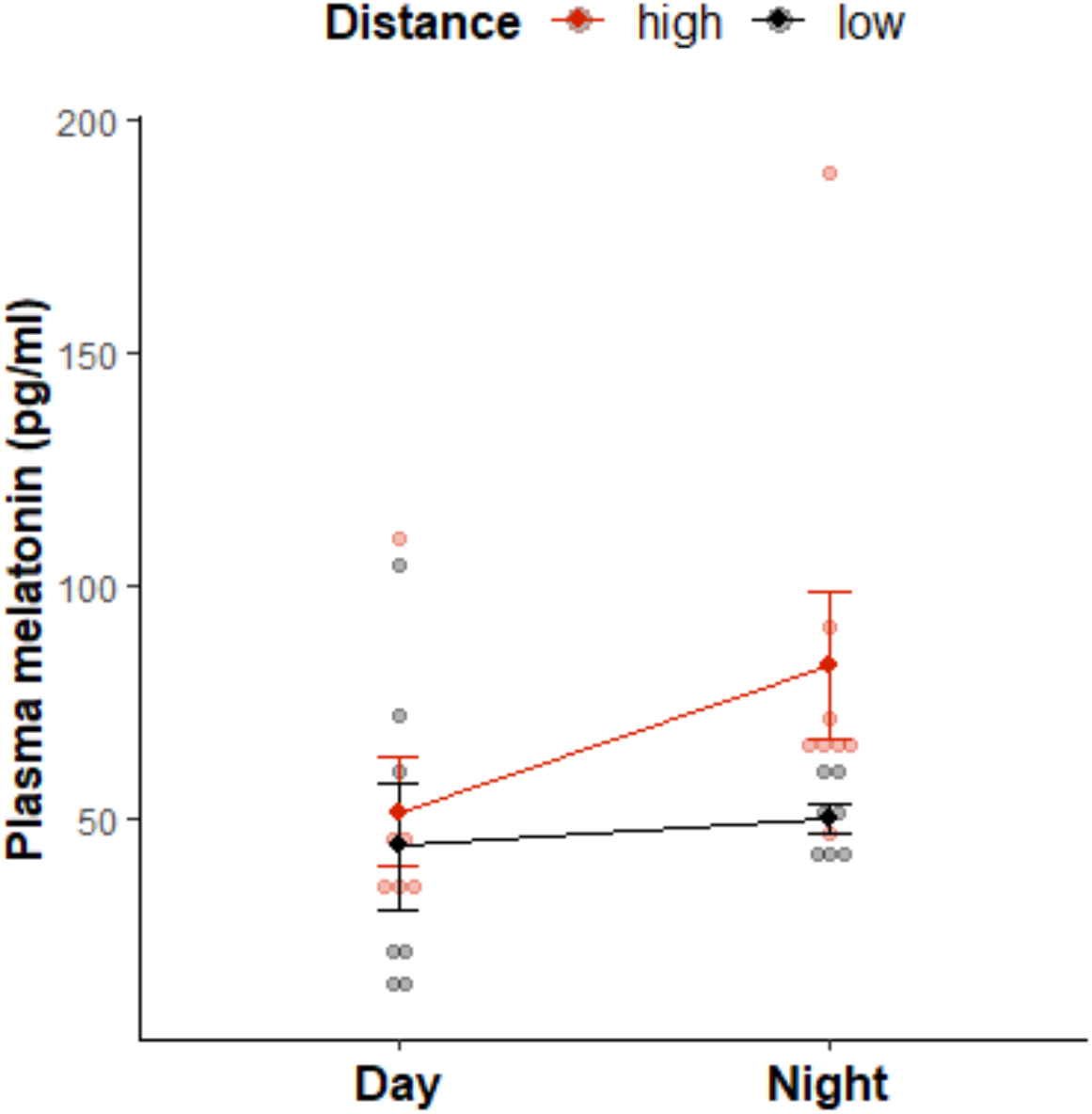
Plasma melatonin levels do not differ significantly between individuals classified as proactive or reactive based on their travelled distance, at night or during the day. Melatonin concentrations determined by EIA in plasma (pg/ml) of threespine sticklebacks collected at the end of day (n=13) or the end of night (n=15), presented separately for individuals exhibiting high (red dots) or low (black dots) activity during the day. Two-way ANOVA with Time of Day and Distance group as variables: Time of Day p = 0.032; Distance p = 0.056; Interaction p = 0.85. Mean ± SE.

**Table 3.** Two-way ANOVA on melatonin levels with Time of Day and Distance as variables. Shapiro-Wilk normality test showed that residuals were not normally distributed. Two-way ANOVA was performed with base-10 log transformed plasma melatonin data and untransformed distance data.

|  | Df | Sum Sq | Mean Sq | F | p-value |
| --- | --- | --- | --- | --- | --- |
| **Time of Day** | 1 | 1.543 | 1.5431 | 5.186 | **0.032** |
| Distance | 1 | 1.085 | 1.0854 | 4.048 | 0.056 |
| Time of Day: Distance | 1 | 0.010 | 0.0102 | 0.038 | 0.847 |
| Residuals | 24 | 6.435 | 0.2681 |  |  |

For time spent in centre, a measure of anxiety, there was a significant interaction between time of day and behaviour type (being in the proactive or reactive group) (Two-way ANOVA, interaction p = 0.005, Table 4). Reactive individuals, which spent most of their time away from the centre of the aquarium, showed significant variation in melatonin between night and day (Tukey multiple comparison test, adjusted p-value = 0.001), while individuals that were characterized as proactive because they spent a larger proportion of time in the centre of the aquarium showed very little variation in melatonin between night and day levels (Tukey multiple comparison test, adjusted p-value = 0.998, figure 4). Melatonin levels differed significantly during the day between the high and low groups since individuals in the group with a low propensity to spend time in the centre had very low melatonin levels at that sampling point (Tukey multiple comparison test, adjusted p-value = 0.018) but this difference was not seen at night (adjusted p-value = 0.686).

**Figure 4.**
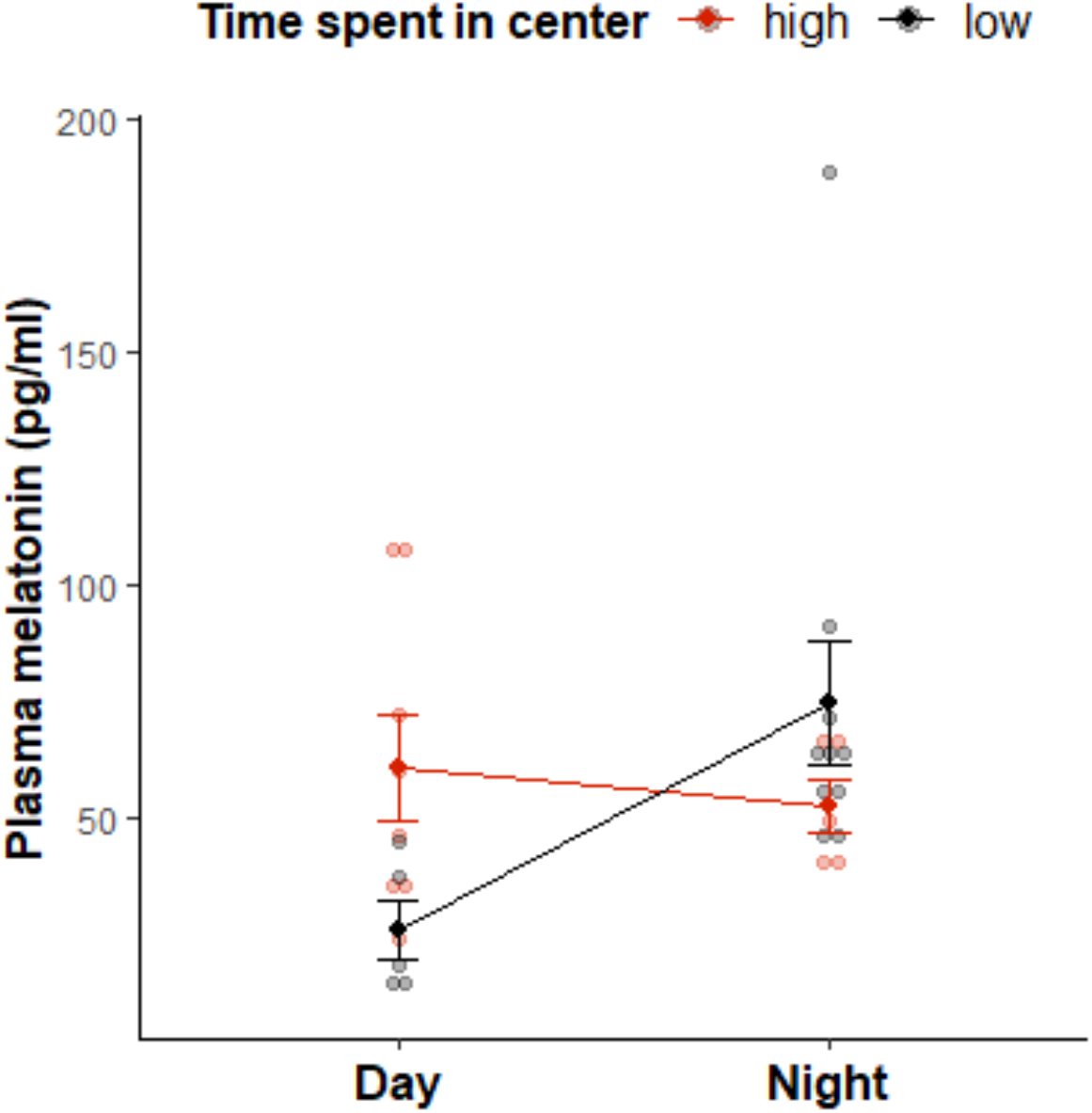
Time of day and behaviour type for time spent in centre showed a significant interaction on plasma melatonin levels. Melatonin concentrations determined by EIA in plasma (pg/ml) of threespine sticklebacks collected during the day (n=13) or night (n=15), presented separately for individuals spending a high proportion (red dots) or a low proportion (black dots) of time in the centre of the aquarium during the day. Two-way ANOVA with Time of Day and Time spent in centre group as variables: Time of Day p = 0.012; Time spent in centre p = 0.164; Interaction p = 0.005. Mean ± SE.

**Table 4.**
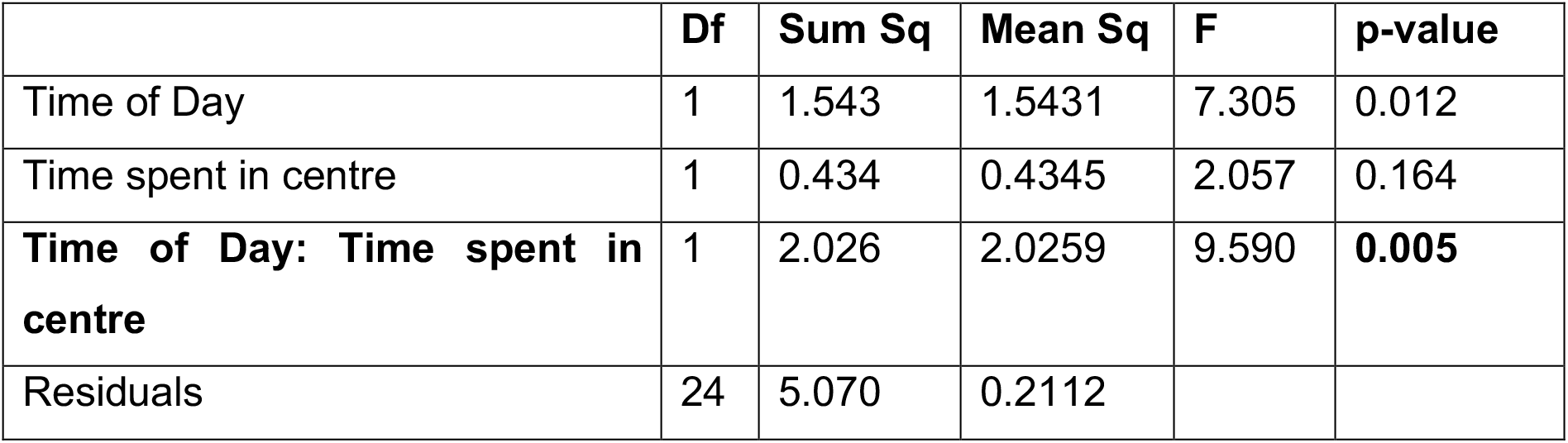
Two-way ANOVA on melatonin levels with Time of Day and Time spent in centre as variables. Shapiro-Wilk normality test showed that residuals were not normally distributed. Two-way ANOVA was performed with base-10 log transformed plasma melatonin data and untransformed time spent in centre data.

Focusing on predator response, we found no significant differences in melatonin levels between individuals split in “high” and “low” group for the behaviours Time to freeze and Time spent frozen (interaction term was not significant, data not shown).

## DISCUSSION

The link between melatonin levels and personality, locomotor activity or stress response is well documented in fish (Azpeleta et al., 2010; Bayarri et al., 2004; López-Olmeda et al., 2006; Sánchez-Vázquez et al., 2019; Tudorache et al., 2018; Vera et al., 2005). In addition to this general association between melatonin levels and behaviour, in zebrafish, a diurnal species, rhythmicity of melatonin secretion is more pronounced in proactive individuals than in reactive ones (Tudorache et al., 2018). Proactive zebrafish (with a faster emergence time in a novel environment) have significantly higher melatonin levels at night than during the day, while this difference is not present in reactive individuals. Here, in threespine sticklebacks, we measured large inter-individual variation in melatonin levels and found that activity was positively correlated with plasma melatonin concentration measured at night. However, contrary to our prediction, we did not observe any significant difference in nigh-day variation in melatonin concentration between very active and less active groups. However, opposite of what we predicted, we found that individuals that spent most of their time away from the centre of the aquarium (classified as reactive) showed large variation in melatonin between night and day, while this rhythm was not seen in proactive individuals that spent a higher proportion of their time in the centre of the aquarium. Our results suggest that interindividual variation in melatonin levels is widespread in wild individuals. The positive association of this variation in melatonin with variation in activity suggests that melatonin may directly modulate activity in threespine sticklebacks, as found in other species. Our results also suggest that the pattern of a difference in melatonin rhythm between proactive and reactive individuals may be widespread in fish, but that whether a species is diurnal or nocturnal could be the cause of differences observed between zebrafish and our own experiment.

### Inter-individual variation in melatonin levels

It had already been shown that melatonin levels are significantly higher at night in the brain of threespine sticklebacks than during the day (Kulczykowska et al., 2017) and our measurements in plasma confirm that observation, while also clearly showing that individual wild threespine sticklebacks vary in their melatonin levels even when placed in stable laboratory conditions for a year. Our measurements show that melatonin level in plasma is variable between individual threespine sticklebacks is in accordance with findings in European sea bass (Bayarri et al., 2002), Senegal sole (Bayarri et al., 2004) and Atlantic salmon (Porter et al., 2001; Randall et al., 1995). This large variation could be the result of the sticklebacks’ environment during early life that affects phenotypes in an irreversible manner (developmental plasticity) or of genetic variation, which could be dissected by studying lab-reared individuals for which the environment can be manipulated. The repeatability of these melatonin measurements for a given individual will also need to be quantified. In threespine sticklebacks, some individuals are more predictable in their behaviours than others (i.e. they have a higher repeatability) and it would be interesting to test if physiological traits such as melatonin levels also show repeatability (Jolles et al., 2019). Interestingly, melatonin, similar to other hormones such as cortisol, is released through gills and has been measured in a non-invasive manner in environmental water of rainbow trout (*Oncorhynchus mykiss*) or European perch (*Perca fluviatilis*) (Brüning et al., 2015; Ellis et al., 2005; Scott and Ellis, 2007) but not in threespine sticklebacks. The use of this method would allow to measure melatonin in the same individual repeatedly.

### Correlation between melatonin levels and activity

In zebrafish, individuals that showed the largest amplitude of change in melatonin levels between day and night were also classified as proactive (exhibiting more risky behaviour) and were more active when daylight appeared (Tudorache et al., 2018). This association between high melatonin levels at night and high activity during the day was measured in unmanipulated animals and contrasts with what is found when melatonin levels are experimentally raised at night in goldfish and sea bass, since in that case a decrease in locomotor activity levels is then observed during the day (Azpeleta et al., 2010; López-Olmeda et al., 2006) (Herrero et al., 2007). Our results are partly in line with what was found in *Danio rerio*, since sticklebacks showed a significant positive association between their activity level measured as the total distance they travelled during daytime and their night time (darkness) melatonin levels. This could reflect a causal relationship, with melatonin absolute concentration at night causing changes in activity the next day. This contrasts with what was found in fish species where melatonin levels were artificially manipulated, potentially because manipulation does not reproduce the physiological context, the moment of maximum levels, nor the duration of melatonin elevated levels. Furthermore, the dose used during a manipulation could also affect the results on behaviour, either because there is a non-linear relationship between melatonin level and behaviour or because of receptor saturation. Finally, manipulation studies in different species do not always have the same findings, suggesting that the link between melatonin and behaviour could be species-specific, or even population or individual-specific. If there is a causal link between melatonin levels and behaviour and the observed variation in melatonin levels results from genetic variation, one could then shed light on an evolutive question: is there an advantage to melatonin and behaviour to be linked? For example, a hypothesis would be that a higher activity level during the day could be advantageous and directly be induced by a more pronounced melatonin rhythm, which would be under selection. It is also possible that the association we quantified between melatonin levels and activity instead reflects behaviour influence on melatonin secretion rather that the other direction. Locomotor activity has already been described as an entraining signal and represents a mechanism for an endogenous behaviour to feedback and influence circadian function (Hughes and Piggins, 2012). Mechanism of the causal link between behaviour and consequent melatonin levels, if it exists, would thus be worth investigating. To further dissect this association between melatonin and activity, it would be useful to simultaneously follow melatonin levels and locomotor activity in threespine sticklebacks over several days. Measuring melatonin in water, as suggested above, would allow to repeat measurements on the same fish, simultaneously monitoring behaviour or physiology while eliminating sampling stress.

It is worth noting that we did not find any correlation between individual predator response and plasma melatonin concentration, in contrast to activity. Both exploration and predator response variables measured in our study are consistent with behavioural studies already carried out in threespine sticklebacks (Aubin-Horth et al., 2012; Berger and Aubin-Horth, 2020; Grécias et al., 2018, 2017; Lacasse and Aubin-Horth, 2012). This suggest that melatonin levels do not affect daylight response to predators.

### Amplitude of melatonin level changes and behaviour

In zebrafish, individuals labelled as proactive because of their fast emerging time in a new environment and their high activity levels during the day present a higher amplitude in melatonin than reactive individuals (Tudorache et al., 2018). We thus predicted in wild threespine sticklebacks that individuals with a pronounced melatonin rhythm would also exhibit high activity levels during the day, but we found no such significant association for this behaviour. However, we found that individuals that spent most of their time away from the centre of the aquarium (classified as reactive) showed large variation in melatonin between night and day, while this rhythm was not seen in proactive individuals that spent a higher proportion of their time in the centre of the aquarium. Thus, in this nocturnal species, it is the reactive individuals that show the most contrasted melatonin rhythm. This suggests that melatonin levels during the day, which are higher in proactive individuals, could be the cause of this low anxiety. Indeed, experimentally raising melatonin levels for 24 hours in zebrafish results in an anxiolytic effect, measured as time spent near the surface, a measure of boldness and low predator response (Genario et al. 2020). If this experimental result applies to sticklebacks, proactive individuals that spend more time in the centre of the tank would exhibit this reduced anxiety behaviour potentially because of their high melatonin levels, an hypothesis which needs to be tested directly in this species. This effect of melatonin on anxiety could explain the opposite pattern of proactive or reactive individuals having the most contrasting daily melatonin rhythm in zebrafish and sticklebacks since they were found for different behaviours. Indeed, measuring exploration using time to emergence in a novel environment (as in zebrafish) tests a different behaviour than measuring time spent in the centre of the tank. Therefore, the opposite pattern found in the two species could both be caused by melatonin but through different molecular and endocrine pathways that result in opposite behavioural effects.

Since activity measured as distance travelled was not correlated with propensity to spend time in the centre, the sticklebacks that are classified as proactive based on activity are not necessarily the same that are classified as proactive when using this anxiety measurement. This suggests that although we originally grouped them into an “exploration” category, they measure different components of behaviour and that action of melatonin on activity and anxiety could thus happen through different pathways and have different effects. Supporting this, in zebrafish, individuals who favour swimming close to the walls are classified as exhibiting high anxiety, since this behaviour can be modulated using anxiolytic drugs targeting the 5HT1A serotonin receptor (Maximino et al., 2013). Experimentally treated individuals spend less time close to the walls and freeze significantly less, but do not show more total swimming activity. Thus, in that case, these behaviours seem to be controlled by different neuroendocrine pathways.

Finally, another explanation for the lack of a relationship between melatonin rhythm amplitude and activity in sticklebacks compared to zebrafish would be that the effects of melatonin levels on activity would be measurable at night in this nocturnal species. Activity measured at night may not be correlated to activity during the day, since a negative correlation between night activity and total daily activity has been shown in sticklebacks held in the laboratory (Brochu and Aubin-Horth 2021).

### Future directions

The main difficulty of our study was to measure for the first time the concentration of melatonin by EIA in small samples: the brain and plasma of threespine sticklebacks. We adapted the manufacturer protocol but the concentration of melatonin in the samples was nevertheless close to the lower limit of detection of the EIA kit, in particular in samples collected during daytime. We did find higher melatonin concentrations in plasma at night as expected but not in the brain. Similar melatonin concentrations have been measured in the brain of threespine sticklebacks using HPLC, but a significant difference was found between melatonin concentration during the day and at night (Kulczykowska et al., 2017). This suggest that our C18 column extraction and/or EIA assay protocol would need to be improved for that tissue. We suggest to improve brain tissue lysis using a sonication probe or silica beads and lysing tissues in methanol instead of phosphate-buffered saline (see Sokołowska et al., 2004). Furthermore, as we were experimenting with a new protocol for this species, we decided to focus on two time points over a 24-hour cycle. It would be appropriate to repeat the same experiment using more numerous sampling time points, as done in zebrafish for example (Tudorache et al., 2018) and over several 24-hour cycles.

Since threespine sticklebacks are nocturnal but the most active individuals tend to be more active during the day than at night (Brochu and Aubin-Horth, 2021; Quinn et al., 2012), following behaviour during one or several complete light-dark cycle or testing behaviours both during day and night could be relevant in future studies. Including other behaviours such as time to emerge in a new environment would offer a more complete picture of the link with melatonin rhythms. Finally, dissecting the proposed link between melatonin secretion and anxiety would necessitate to control melatonin levels and directly test anxiety-related behaviours such as wall-hugging.

### Conclusion

Threespine stickleback is a model for behavioural biology (Huntingford and Ruiz-Gomez, 2009) but also for the study of host-parasite relationships (Barber and Scharsack, 2010). *Schistocephalus solidus* infection influence threespine sticklebacks’ behaviour and physiology (Berger et al., 2021; Berger and Aubin-Horth, 2020; Grécias et al., 2018, 2017; Hébert et al., 2017; Øverli et al., 2001; Quinn et al., 2012; Scharsack et al., 2007; Talarico et al., 2017). Serotoninergic turnover is decreased in the brain of infected fish (Øverli et al., 2001) and serotoninergic activity is involved in the control of locomotor activity (Winberg et al., 1993). It is thus possible to predict that *Schistocephalus solidus* could alter melatonin rhythms in the host, resulting in behaviour alteration and potentially manipulation, as suggested in Nile tilapia (Ellison et al., 2018), in human (Claustrat et al., 1998) and in mice (Rijo-Ferreira et al., 2018). Apart from parasitic infection, the dysregulation of melatonin secretion is also involved in different human diseases (Karasek, 2004; Pacchierotti et al., 2001; Pandi-Perumal et al., 2017; Reiter et al., 2007). Thus, understanding the melatonin system in animal models can contribute to a better understanding of human physiology and disease and in vertebrates in general. Biomarkers of biological functions in threespine sticklebacks are used as an indicator of water contamination (Sanchez et al., 2008). Melatonin is already described to be protective against some chemicals (Asghari et al., 2017) or to be modified by others (Kulczykowska et al., 2004). Melatonin secretion in sticklebacks could thus be considered as a candidate biomarker in ecotoxicology and inter-individual variation in this hormone thus must be monitored and included in the analysis. Overall, our study suggests that melatonin may directly modulate activity and anxiety-associated behaviours in wild sticklebacks, and that interindividual variation in melatonin rhythm is widespread in wild individuals but with different pattern of variation depending on the species.

## ACKNOWLEDGMENTS

We thank Verônica A. Alves and Marie-Pier Brochu for their help with field sampling, tissue sampling and lab experiments and the personnel of the Laboratoire Aquatique de Recherche en Sciences Environnementales et Médicales (LARSEM) at Université Laval for their help with fish rearing.

